# OmniClustify^XMBD^: Uncover putative cell states within multiple single-cell omics datasets

**DOI:** 10.1101/2023.12.22.573159

**Authors:** Fan Yang, Yitao Zhou, Feng Zeng

## Abstract

Clustering plays a pivotal role in characterizing cell states in single-cell omics data. Nonetheless, there is a noticeable gap in clustering algorithms tailored for unveiling putative cell states across datasets containing samples with diverse phenotypes. To bridge this gap, we implement an innovative method termed OmniClustify^XMBD^, which integrates adaptive signal isolation with cell clustering. The adaptive signal isolation effectively disentangles gene expression variations linked to distinct factors within individual cells. This separation restores cells to their inherent states, free from external influences. Concurrently, a clustering algorithm built upon a deep variational Gaussian mixture model is devised to identify these putative cell states. Experiments showcase the effectiveness of OmniClustify^XMBD^ in identifying putative cell states while minimizing the influence of various undesired variations, including batch effects and random inter-sample differences. Moreover, OmniClustify^XMBD^ demonstrates robustness in its results across different clustering parameters.

## 1 Introduction

The rapid evolution of single-cell omics has catalyzed a transformative paradigm shift within the life sciences. In recent years, single-cell omics has yielded ground-breaking achievements in biomedical research, ranging from elucidating the mechanisms governing cell fate decisions during development to dissecting aberrant transcriptomic alterations in disease contexts. However, with the expanding repository of single-cell data, it is becoming increasingly evident that the complexities and challenges associated with clustering diverse data encompassing diverse samples and conditions are on the rise. These inherent undesired variations across samples and conditions introduce additional intricacies that hamper the efficacy of clustering methods in addressing these complex scenarios.

Desipte the numerous clustering algorithms, only a select few aim to eliminate the influence of these undesired variations and obtain consensus clustering across datasets [1]. DESC is one such pioneering method, addressing clustering and batch correction through a two-step procedure involving auto-encoder pre-training and deep clustering [2]. However, due to the separation of these steps, the performance of DESC remains suboptimal. On the other hand, scDML leverages initial clustering to identify cell triplets [3] and utilizes contrastive learning to refine the clustering and eliminate batch effects [4]. However, the initial clustering may generate incorrect cell triplets, potentially affecting the final clustering results.

We introduce OmniClustify^XMBD^, a method designed to uncover putative cell states with robust clustering while mitigating undesired variations. Our method combines adaptive signal isolation with deep variational Gaussian-mixture clustering. This involves iterative process aimed at estimating and attenuating residual variations linked to distinct factors in the remaining data. Simultaneously, clustering is applied on the remaining data to characterize putative cell states. To the best of our knowledge, OmniClustify^XMBD^ represents the first approach to seamlessly integrate these two distinct tasks, achieving accurate clustering and effective batch correction concurrently.

## 2 Method and Material

### 2.1 OmniClustify^XMBD^

OmniClustify^XMBD^ consists of two essential components. The first component is meticulously designed to isolate the multifaceted influences stemming from diverse factors acting upon individual cells. Once these influences are effectively isolated, the remaining gene expression signals encapsulate the inherent cell states. The second component is strategically engineered to execute the clustering of cells predicted on on these refined gene expression signals. Notably, these components are seamlessly interwoven within the framework of deep random-effects modeling, facilitating their simultaneous training (see Figs. 1A and B).

**Fig. 1.**
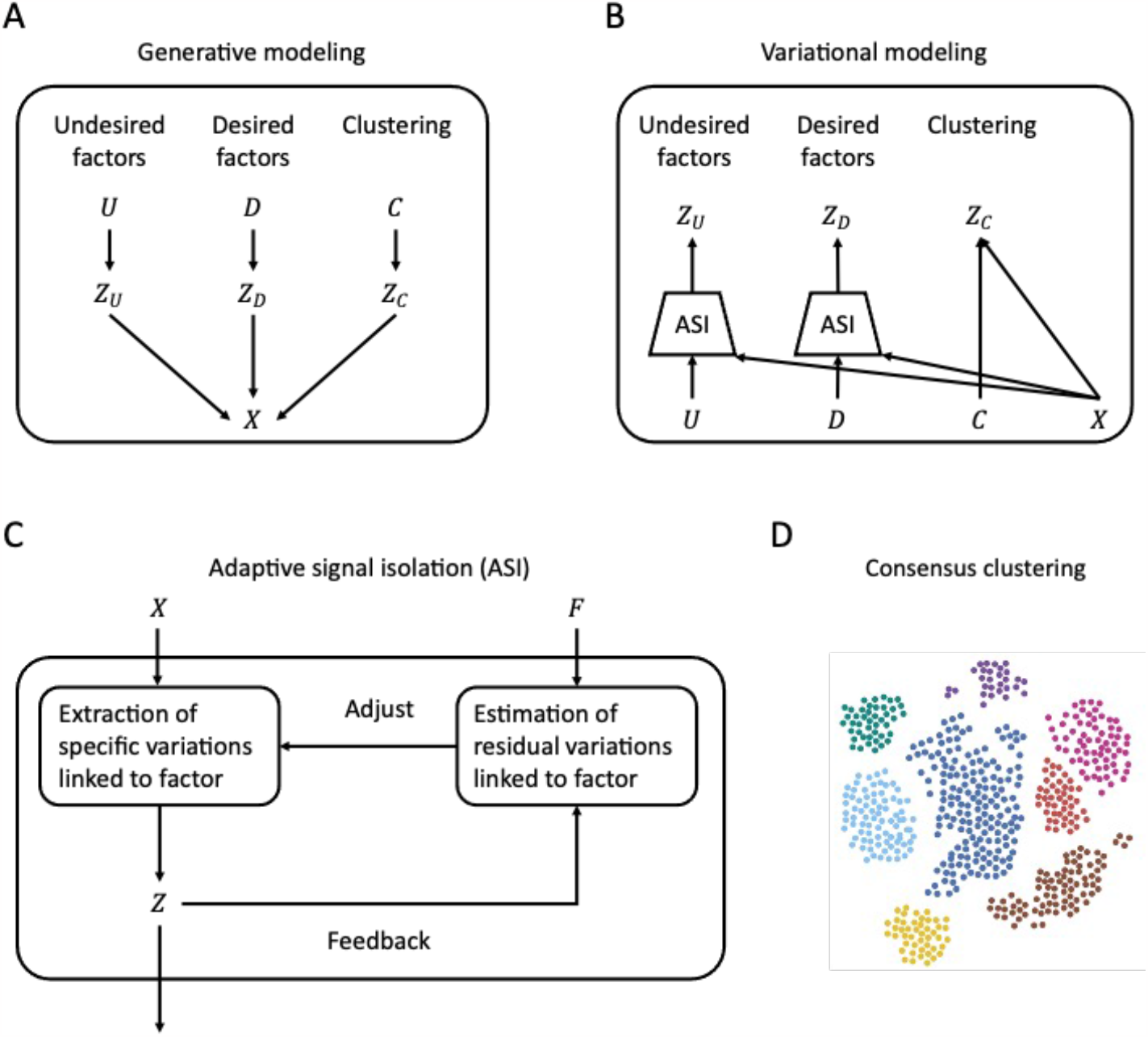
The framework of OmniClustify^XMBD^. **A**, the generation procedure controlled by distinct factors. **B**, the variational inference process for these factors. **C**, the principle of adaptive signal isolation. **D**, the application of OmniClustify^XMBD^ in characterizing putative cell states across various datasets.

### 2.2 Adaptive signal isolation

Single-cell omics data and the auditory perception of sound signals share striking similarities. Both encompass a primary signal, referred to as the target signal. In the context of single-cell omics, this target signal pertains to actual gene expression, whereas in auditory perception, it encapsulates meaningful information. Furthermore, the signals received in both domains are accompanied by diverse superimposed noise. The sources and characteristics of this noise vary significantly, rendering accurate preemptive modeling a formidable challenge. The profound challenge, shared by these ostensibly distinct disciplines, resides in the effective elimination of this noise while preserving the fidelity of the target signal.

The conceptual roots of this endeavor can be traced back to the seminal contributions of Norbert Wiener and Rudolf E. Kalman, who laid the foundation for optimal filtering [5, 6]. Today, this pursuit is commonly recognized as active noise control, a concept that has found successful application across diverse engineering domains, spanning communication, speech processing, and biomedical signal analysis.

Taking inspiration from this concept, we have devised an adaptive algorithm for the isolation of distinct variations within single-cell omics data (see Fig 1C). Adaptive signal isolation consists of two essential components. One is to extract variations belonging to specific factors. The other is to estimate the remaining information needed further isolation.

We implemented these processes through a random-effects modeling approach [7]. The factors present in single-cell omics data are categorized into two types: undesired variations, denoted as a binary vector *U* (e.g., batch variation and random inter-individual differences), and desired variations, denoted as *D* (e.g., phenotypic variations related to diseases or perturbations). For illustrative purpose, let’s exclusively consider the former—*U*. In traditional latent variable models, *U* is specified to represent a cell state prototype for cells, collectively defining the landscape of cellular variations influenced by *U*. However, the variance *δ*_*U*_ related to *U* is often inadequately modeled, despite its significance in quantifying the precision of information extraction. This variance characterizes the degree of deviation exhibited by each cell from its prototype and, consequently, captures the random effects reflecting the influence of variations allocated to *U*. The inclusion of this precision parameter becomes crucial since unrelated variations in *U* could inadvertently introduce superfluous and irrelevant information to the isolated signal, inflating cell heterogeneity,.

First, let’s consider the modeling of *U* to address this issue. Specifically, in the first step, the generation of *U* is modeled by using a combination of beta-binomial distributions:

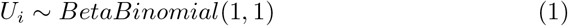

For *δ*_*U*_, we assume that it is from a standard normal distribution:

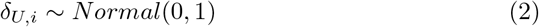

Once *U* and *δ*_*U*_ are specified, the latent variable *Z*_*U*_ and its variance 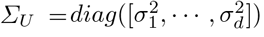, where *d* is the dimension of latent variable *Z*, are generated through a neural network function *ϕ*_*G*_(*U, δ*_*U*_) as follows:

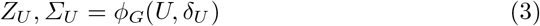

Similarly, the latent variable *Z*_*D*_ and its variance *Σ*_*D*_ are generated using a similar procedure and another neural network function *ψ*_*G*_(*D, δ*_*D*_) when the prototype *D* and the deviation *δ*_*D*_ are provided.

To estimate the parameters of the generative model described above, we employ variational approximation techniques [8]. For instance, when dealing with the undesired factor *U*, we establish the following variational model:

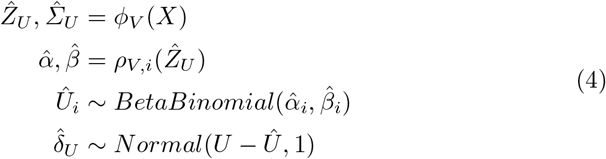

In this model, *ρ*_*V*_ () denotes a neural network function, and a similar approach is applied to the desired factor *D*. It is worth noting that we rely on the difference *U* − *Û* to estimate the deviation 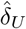. A small value of *U* − *Û* indicates that the algorithm has effectively extracted the related variations, while a substantial value of *U* − *Û* will prompt the algorithm to adapt its parameters to optimize the extraction process.

### 2.3 Deep variational Gaussian mixture modeling

To cluster cells, we adopt the framework of deep variational Gaussian mixture modeling proposed in scVAE [9]. This framework can be seamlessly incorporated into the above deep random-effects modeling. Briefly, when the cluster label *C* of a cell is specified, its latent variable *Z*_*C*_ and its variance *Σ*_*C*_ is generated using the following generation process:

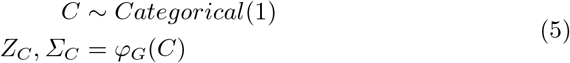

Its accompanying variational model is as follows:

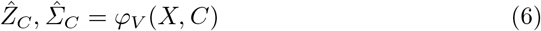

The collection of the latent variables *Z*_*U*_, *Z*_*D*_, and *Z*_*C*_ characterizes the complete state of a cell. They are used to generate its gene expression signal through the zero-inflated negative binomial (ZINB) distribution.

### 2.4 Code variability

OmniClustify^XMBD^ and materials are available for academic use on GitHub at https://github.com/ZengFLab/OmniClustify_XMBD.

## 3 Results

### 3.1 An example of concurrent signal isolation and cell clustering

To demonstrate the adaptive capabilities of OmniClustify^XMBD^ in selectively isolating the signals attributed to distinct factors while preserving the remaining signal and elucidating putative cell states, we utilized a single-cell RNA sequencing (scRNA-seq) dataset derived from 41,650 cells originating from four distinct colon regions in five healthy males [10]. This dataset inherently displays a variety of variations, comprising factors of interest (see Figs. 2A-C). Specifically, it encompasses variations signifying individual identity, which, although informative, are undesired in the context of our analysis. Moreover, it encapsulates variations associated with colon regions, which are of particular interesting desired when investigating cell heterogeneity influenced by local colon environments, i.e., microbial compositions. Most notably, the predominant source of variation in this dataset is attributed to distinct cell types.

**Fig. 2.**
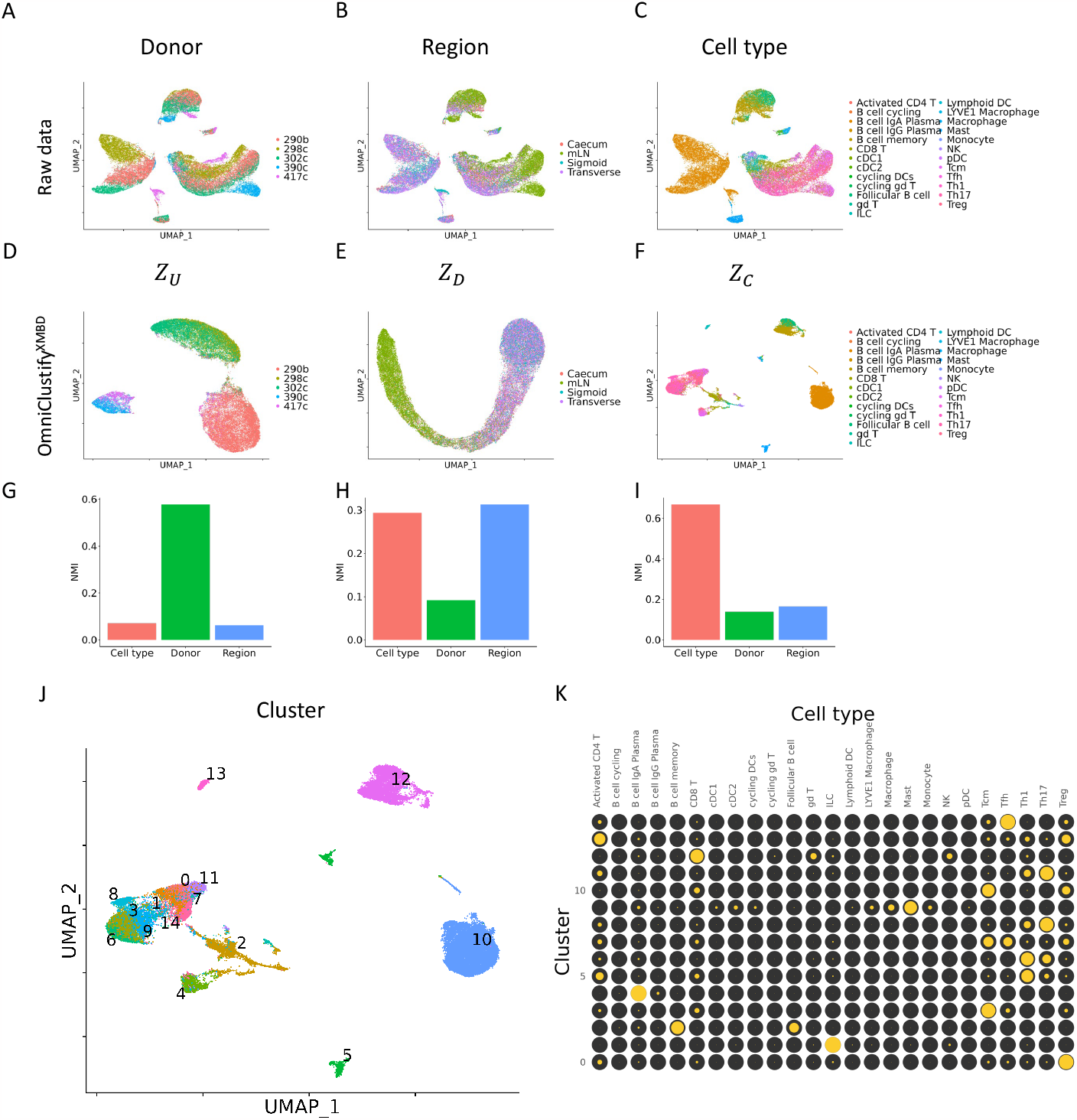
Illustration of concurrent signal isolation and cell clustering. **A-C**, UMAP plots of raw data colored by donor identity, colon region, and cell type, respectively. **D-F**, UMAP plots for the decomposition components *Z*_*U*_, *Z*_*D*_, and *Z*_*C*_. **G-I**, NMI metrics quantify the extent of information captured by distinct decomposition components. **J**, visualization of OmniClustify^XMBD^ clustering. **K**, the correspondences between cell clusters and cell type annotations, highlighted by yellow dots, with dot size indicating the proportion of cells within a cluster mapped to the specific cell type.

In this experiment, we leveraged OmniClustify^XMBD^ to deconvolute the complexities within this dataset. Importantly, our experimental approach deliberately omitted the use of cell type annotation information, with the explicit anticipation that OmniClustify^XMBD^ would autonomously discern and identify these distinct cell states. As shown in Figs. 2D-F, we calculated the two-dimensional uniform manifold approximation and projections for the decomposition components *U, D*, and *C*, respectively. These results corroborated that the inherent variations in this dataset were accurately distinguished and separated from one another. The utilization of the normalized mutual information (NMI) index as a metric reveals the extent to which the decomposition components captured related variations (see Figs. 2G-I). The information residing in the latent space of *U* is primarily associated with donors and exhibits limited relevance to regions and individuals. The results of *C* demonstrate a similar trend. Interestingly, alongside the information pertaining to regions, the isolated variations related to *D* unveils insights into cell types, suggesting that local colon environments may exert a substantial influence on the regulation of cell fates.

To validate the clustering results produced by OmniClustify^XMBD^, we conducted a comparison with cell type annotations. The outcomes reveal a high degree of concordance between cell clustering and annotated cell types (see Fig. 2K). This result is exceptionally encouraging, given the entirely unsupervised nature of the analysis. Furthermore, it marks the pioneering instance in which distinct sources of variation within single-cell omics data have been identified and isolated without bias.

### 3.2 Evaluation of cell clustering for single-cell omics

In order to comprehensively assess the efficacy of OmniClustify^XMBD^, we conducted an in-depth evaluation employing gene expression profiles derived from datasets generated during the NeurIPS 2021 Challenge [11]. These datasets encompassed samples obtained from nine healthy individuals, wherein bone marrow mononuclear cell suspensions were distributed to four distinct locations, followed by single-cell sequencing procedures.

The resulting dataset was characterized by a myriad of intricate variations associated with several influencing factors, notably individual identity, facility location, and cell type. To rectify the undesired variations introduced by inter-individual and inter-location differences and to delineate putative cell states within this complex dataset, we harnessed the capabilities of OmniClustify^XMBD^. In our comparative analysis, we also included two established methods, DESC and scDML, both renowned for their dual functionalities encompassing batch effect correction and cell clustering.

As presented in Fig 3, the results of our investigation provide a comprehensive evaluation of OmniClustify^XMBD^’s performance in the context of complex single-cell omics data derived from diverse individuals and facility locations. These findings also offer a benchmark for comparing the effectiveness of OmniClustify^XMBD^ with DESC and scDML, enhancing our understanding of its utility in handling intricate real-world datasets.

**Fig. 3.**
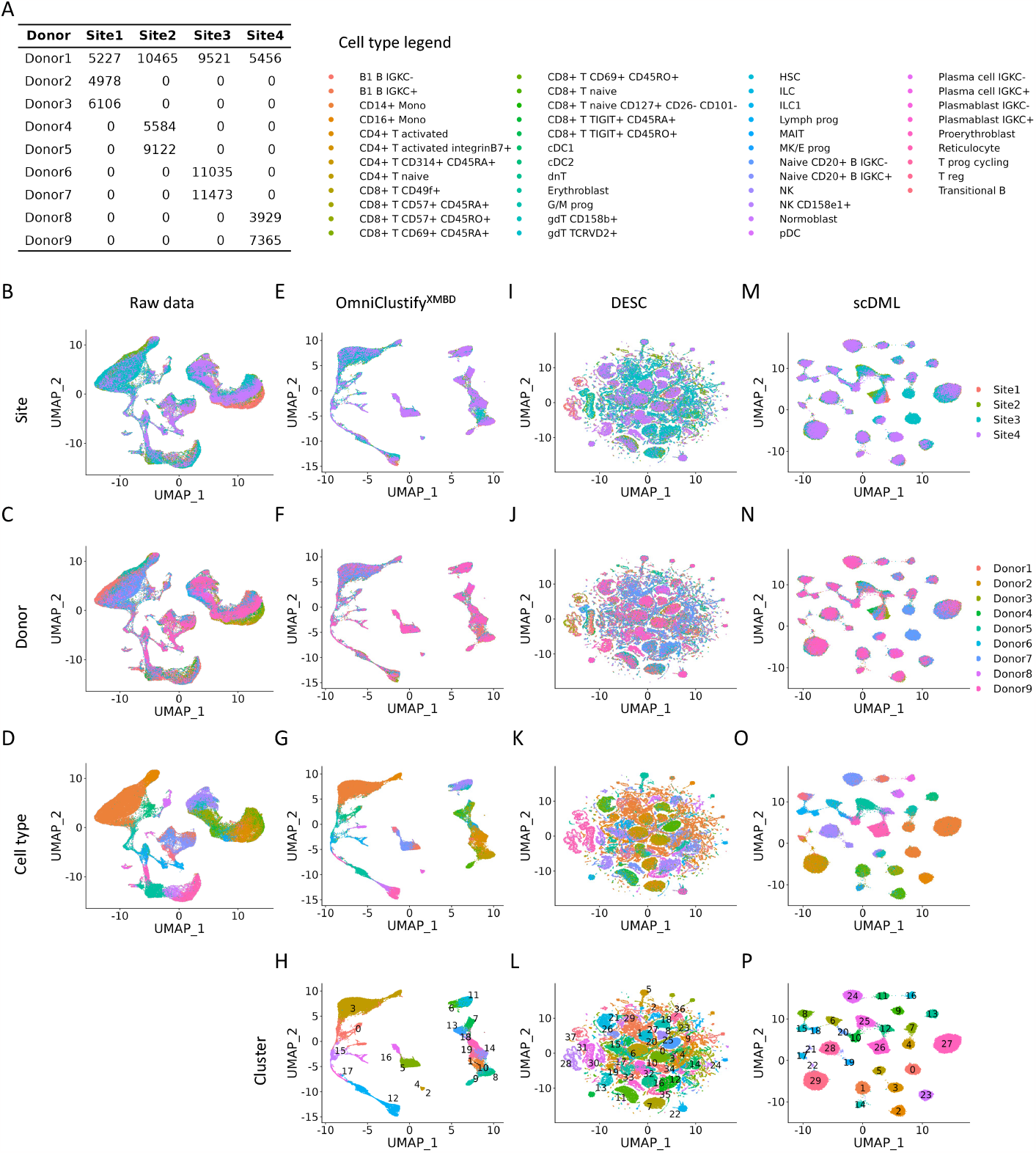
Comparison of cell clustering. **A**, the statistics of the datasets used for the assessment. The datasets were generated by four distinct facilities. **B-D**, UMAP plots of the raw data colored by facility location, donor identity, and cell type, respectively. **E-H**, UMAP plots of *Z*_*C*_ generated by OmniClustify^XMBD^, colored by facility location, donor identity, cell type, and cluster, respectively. **I-L**, the results of DESC. **M-P**, the results of scDML.

### 3.3 Clustering robustness

In order to assess the robustness of the algorithms concerning the configuration of the clustering parameters, i.e., the number of clusters *k*, we conducted an investigation employing a data created by combing two individual datasets derived from different versions of 10x single-cell protocols [12, 13]. One dataset consisted of cells originating from the Jurkat and Raji cell lines, while the other exclusively contained cells derived from the Jurkat cell line.

We systematically varied the number of clusters, setting it to 2, 5, 8, and executed the clustering algorithms OmniClustify^XMBD^, DESC, and scDML for each configuration. The results are presented in Fig. 4. Visual inspection of the results revealed that OmniClustify^XMBD^ consistently delivered stable clustering results. In contrast, both DESC and scDML exhibited notable sensitivity to the chosen clustering parameter, resulting in clustering outcomes that exhibited significant disparities.

**Fig. 4.**
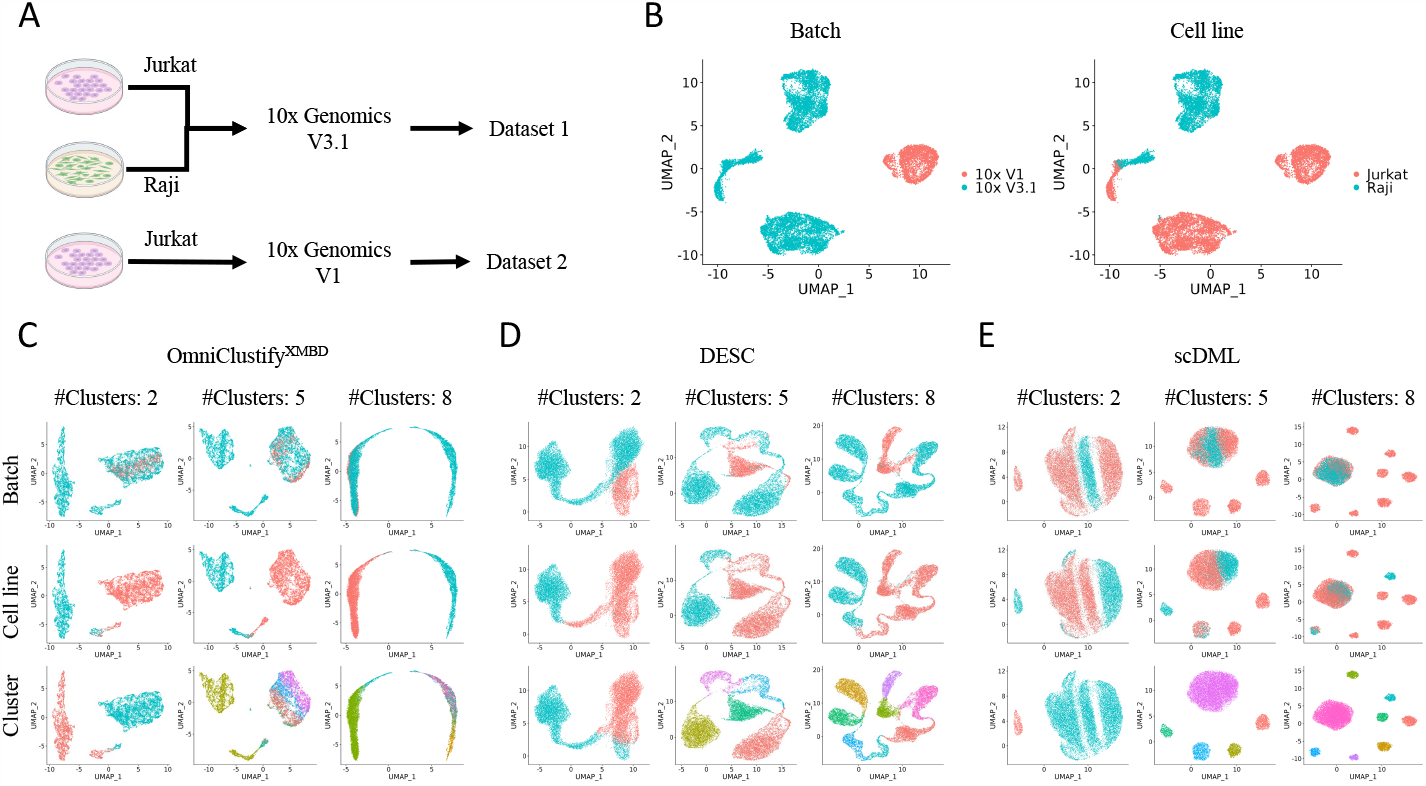
Evaluating the ability of algorithms in producing robust clusterings. **A**, the creation of the testing dataset. **B**, the inherent variations in this data. **C**, the results of OmniClustify^XMBD^ when choosing the parameter of cluster number from the set {2,5,8}. **D**, the results of DESC. **E**, the results of scDML.

## 4 Discussion

We have presented OmniClustify^XMBD^, an innovative clustering methodology specifically designed for the identification of putative cell states within the intricate landscape of diverse single-cell omics datasets. This method represents an extension of our recent work in adaptive signal isolation [14, 15], now coupled with variational Gaussian mixture modeling. OmniClustify^XMBD^ stands as an exceptional advancement in its capacity to robustly and accurately delineate distinct variations while generating dependable cell clusterings. It emerges as an invaluable asset, particularly in the context of personalized medicine research, where comprehensive insights gleaned from large-scale single-cell omics atlases are of paramount significance.

## 5 Fundings

This work has been supported in part by the National Natural Science Foundation of China (61503314, to F.Z.), Natural Science Foundation of Fujian Province (2019J01041, to F. Z.), and General Project of the Xiamen Natural Science Foundation (3502Z20227180, to F.Y.).

## Notes

### Competing Interest Statement

The authors have declared no competing interest.

